# Enhancing ML-based binder design with high-throughput screening: a comparison of mRNA and yeast display technologies

**DOI:** 10.64898/2026.02.12.705611

**Authors:** Zhiyuan Yao, McGuire Metts, Avery K. Huber, Jingjing Li, Tomoaki Kinjo, Henry Dieckhaus, Amrita Nallathambi, Albert A. Bowers, Brian Kuhlman

## Abstract

Recent advances in machine learning (ML)-based protein design methods have enabled the rapid *in silico* generation of large libraries of miniprotein binders with minimal manual input. While computational design capacity has scaled rapidly, experimental validation methods have lagged, creating a bottleneck in binder discovery pipelines. Here, we apply mRNA display to screen an ML-designed miniprotein binder library and directly compare its performance with the more widely used yeast surface display platform using a single shared DNA library. We screened 2,009 designs targeting the platelet receptor TLT-1 and 3,159 designs targeting the immune receptor B7-H3 across both platforms. While both selection methods reliably identified functional binders, we found that mRNA display preferentially enriched binders with slower dissociation rates. In addition, mRNA display achieved higher library coverage than yeast display, likely rescuing functional designs that are penalized in a cell-based expression system. Biophysical characterization of selected binders from both platforms revealed strong binding affinities and high thermal stabilities. These results showcase the power of integrating ML-based computational design tools with rapid *in vitro* selection technologies, providing a scalable framework for therapeutic miniprotein discovery.

**IMPORTANCE:** Miniprotein binders offer major advantages as next-generation therapeutics, including small size, high stability, and efficient production. In this work, we conduct a side-by-side comparison of mRNA and yeast display as platforms for high-throughput evaluation of *de novo* miniprotein binders. The binders generated here serve as starting points for therapeutics targeting TLT-1 or B7-H3, two clinically relevant molecules.

## INTRODUCTION

Recent advances in machine learning (ML) have led to dramatic breakthroughs in protein design, including structure prediction, sequence generation, and scaffold design ^1^. Tools such as AlphaFold2 ^2^, ProteinMPNN ^3^, and RFdiffusion ^4^ have transformed *de novo* design, enabling the generation of tens of thousands of protein sequences with minimal manual input. In particular, ML-based pipelines have shown remarkable success in designing miniprotein binders — single-domain proteins typically 40-80 amino acids in length — that achieve high affinity and specificity across diverse therapeutic targets ^5–8^. These developments highlight the potential of ML-designed miniproteins as next-generation therapeutic candidates.

While computational design capacity has scaled rapidly, experimental validation methods have lagged, creating a bottleneck in the binder discovery pipeline. Many miniprotein design studies rely on low-throughput workflows, manually selecting and expressing only 10–20 candidates for validation ^5–7^. This approach is insufficient to keep pace with the scale of modern ML design and limits our ability to sample the full sequence space or analyze large-scale sequence-function relationships. This gap is particularly problematic for low-success-rate design targets.

Benchmark studies have shown that binder design success varies widely between targets, with success rates for some challenging systems falling below 0.01% ^9^. In such cases, screening limited numbers of sequences is unlikely to recover any true binders. Additionally, limited throughput impairs our ability to systematically evaluate the predictive power of computational scoring metrics or interpret design failures. Although some efforts are underway to benchmark ML-guided design ^9–11^, more datasets are needed to link sequence, structure, and function across large design libraries.

To address this bottleneck, we employ high-throughput screening using mRNA display, a cell-free method commonly used for peptide discovery but not widely adopted for miniprotein binder design. While mRNA display has been previously used to assess properties such as thermostability of designed proteins ^12^, cell-based platforms, particularly yeast display, remain the dominant high-throughput methods for screening binding activity of miniproteins. Yeast display typically requires multiple rounds of cell culture and fluorescence-activated cell sorting ^13^, leading to longer selection cycles and increased hands-on labor. In contrast, mRNA display enables rapid, fully *in vitro* selection, potentially offering shorter turnaround time and lower experimental burden. Here, we perform a direct side-by-side comparison of mRNA and yeast display platforms to assess each platform’s strengths, limitations, and selection biases.

As targets, we selected B7-H3 (also known as CD276), a crucial immune checkpoint protein overexpressed in many cancers ^14^, and TLT-1, a platelet-specific protein involved in blood clotting ^15^. Using a shared DNA library, we evaluate both mRNA and yeast display for screening designed binders, investigate selection biases across platforms, and assess the predictive value of computational metrics. We demonstrate that combining ML-driven design with mRNA display enables the efficient identification of high-affinity miniprotein binders in a single design cycle. This pipeline offers a scalable and efficient framework for accelerating therapeutic miniprotein development.

## RESULTS

### Computational design of miniprotein binders

To initiate the discovery of *de novo* miniprotein binders, we selected two targets: human B7-H3, an immune checkpoint molecule; and human TLT-1, a platelet receptor. Both targets are members of the Ig-like receptor family. For B7-H3, we used a high-confidence model of the human B7-H3 IgV2 extracellular domain (residues 246-357 of UniProt sequence Q5ZPR3) as our target protein in design, which was informed by the solved structure of mouse B7-H3 (92% sequence identity) ^16^. For TLT-1, we used the crystal structure of the human extracellular domain (PDB 2FRG) ^17^ (**Figure 1A**). The target ectodomains were expressed with a C-terminal AviTag^TM^ and enzymatically biotinylated to enable downstream screening.

**Figure 1.**
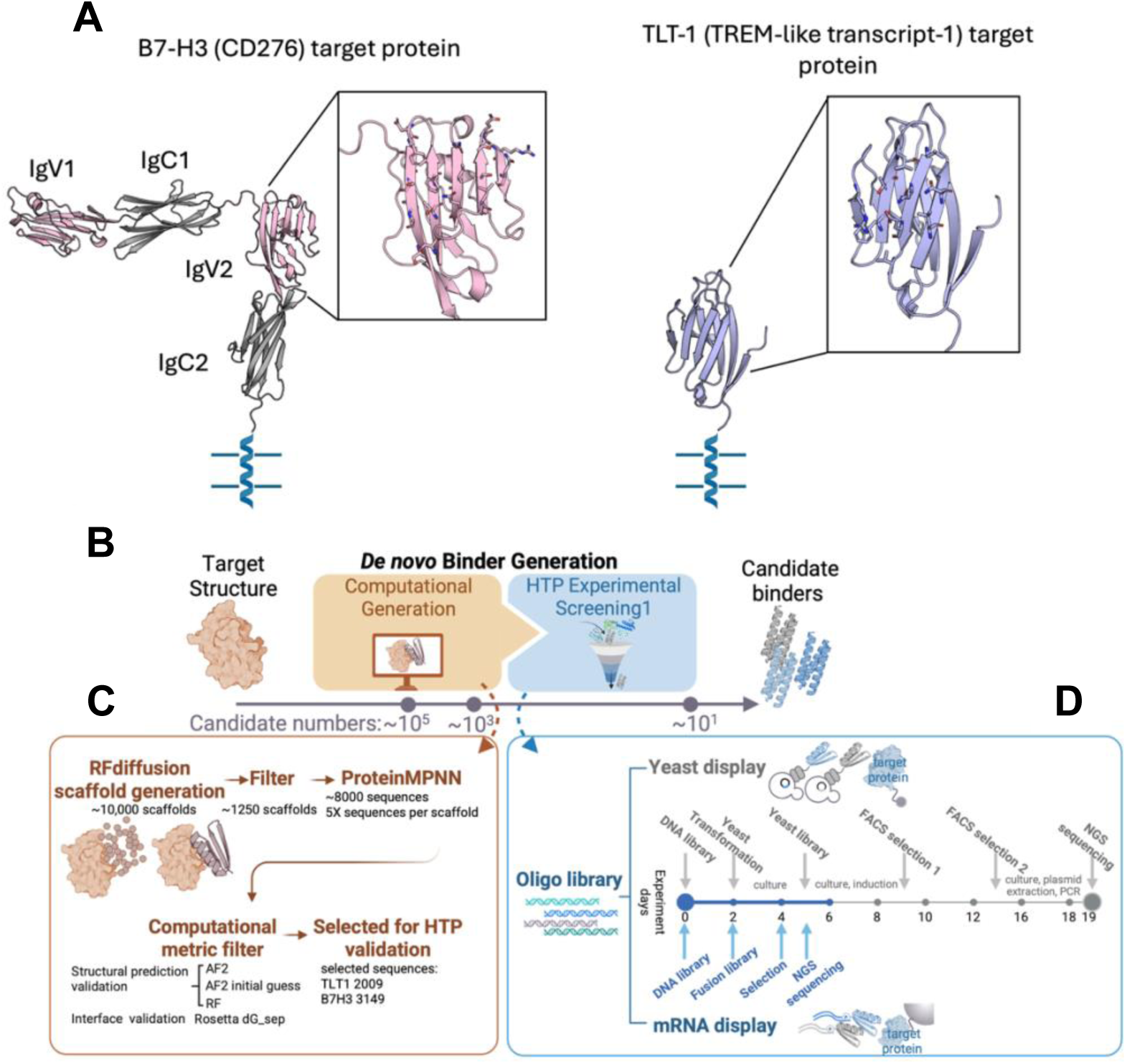
Overview of the computational design and high-throughput screening pipeline for miniprotein binder discovery. (A) Alphafold2 model of the extracellular domain of B7-H3 (left) and a crystal structure of TLT-1 (PDB ID 2FRG) (right) were used as targets for miniprotein binder design. (B) Schematic workflow for miniprotein development between computational design and high-throughput experimental screening. (C) Detailed breakdown of scaffold generation and filtering in computational design phase. RFdiffusion produced ∼10,000 backbones per target, which were filtered to ∼1,250 scaffolds based on geometry and target interface contact. ProteinMPNN sequence design yielded ∼8,000 total candidates. (D) Parallel experimental setup using yeast display (top) and mRNA display (bottom). Both platforms started from a single shared oligo pool, including positive and negative controls.

Using RFdiffusion, we generated an initial pool of around 10,000 miniprotein binder scaffolds against each target structure. Scaffold filtering was applied based on radius of gyration and interface contacts with the target protein. For each target, around 1,250 models were selected for downstream modeling. Sequences were designed for each binder using ProteinMPNN. 5-7 sequences were generated per scaffold, resulting in ∼8,000 designed miniprotein binder sequences. (**Figure 1B, C**).

Designed sequences were evaluated using three independent structural prediction methods: AlphaFold2 (AF2) ^2^, AlphaFold2 initial guess ^9^, and RoseTTAFold (RF) ^18^. These predictions were evaluated using a multi-metric framework incorporating predicted local distance difference test (pLDDT) scores to assess structural confidence of the designed binder, interface predicted aligned error (PAE), and root mean square deviation (RMSD) to the scaffold pose to evaluate binder-target interaction quality. (**Figure S1A, B**). Interface evaluation metrics showed variability across prediction methods (**Figure S1C, D**), emphasizing the importance of consensus-based filtering. Designs that passed thresholds for multiple metrics were selected for experimental validation. Notably, the metric distributions remained continuous after filtering, preserving diversity compared to the unfiltered population (**Figure S1E, F**).

The final experimental library consisted of 2,009 TLT-1-targeting and 3,149 B7-H3-targeting sequences (**Figure 1C**). To assess design diversity, we projected all selected sequences using ESMFold embeddings ^19^ into a 2D t-SNE plot (**Figure S2**). These projections revealed that TLT-1 designs predominantly featured β-strand interactions with the target, while B7-H3 designs favored compact α-helical bundles.

### Parallel high-throughput screening using mRNA and yeast display

To evaluate the efficiency and complementarity of high-throughput binder discovery methods, we performed parallel screening using both mRNA display and yeast surface display, starting from the same oligonucleotide library. The shared DNA library included 2,009 miniprotein sequences targeting TLT-1, 3,149 targeting B7-H3, and 10 PD-1-targeting sequences previously validated as controls ^6^ (**Figure S6**).

mRNA display offers a faster workflow with lower experimental burden, completing selection and sequencing in approximately 6 days, compared to ∼19 days required for yeast display (**Figure 1D**). mRNA display uses cell-free *in vitro* transcription/translation and direct mRNA/cDNA-protein fusion during library selection. In contrast, yeast display involves multiple rounds of cell culture, induction, and FACS sorting. After selection, libraries from both methods were subjected to next-generation sequencing (NGS). Enrichment fold-change (FC) was calculated as follows: for mRNA display, FC was normalized to expression levels (calculated after HA tag purification); for yeast display, FC was calculated relative to the initial input library.

Manhattan plots (**Figure 2A, B**) visualize enrichment scores across all designs for each method. A log₂ fold-change threshold of > 3.5 (i.e., FC > ∼11.3) was used to define high-confidence hits. A subset of enriched candidates from both methods were expressed in *E. coli* (**Figure S7**) and validated using SPR binding assays, with validated hits annotated on the plots.

**Figure 2.**
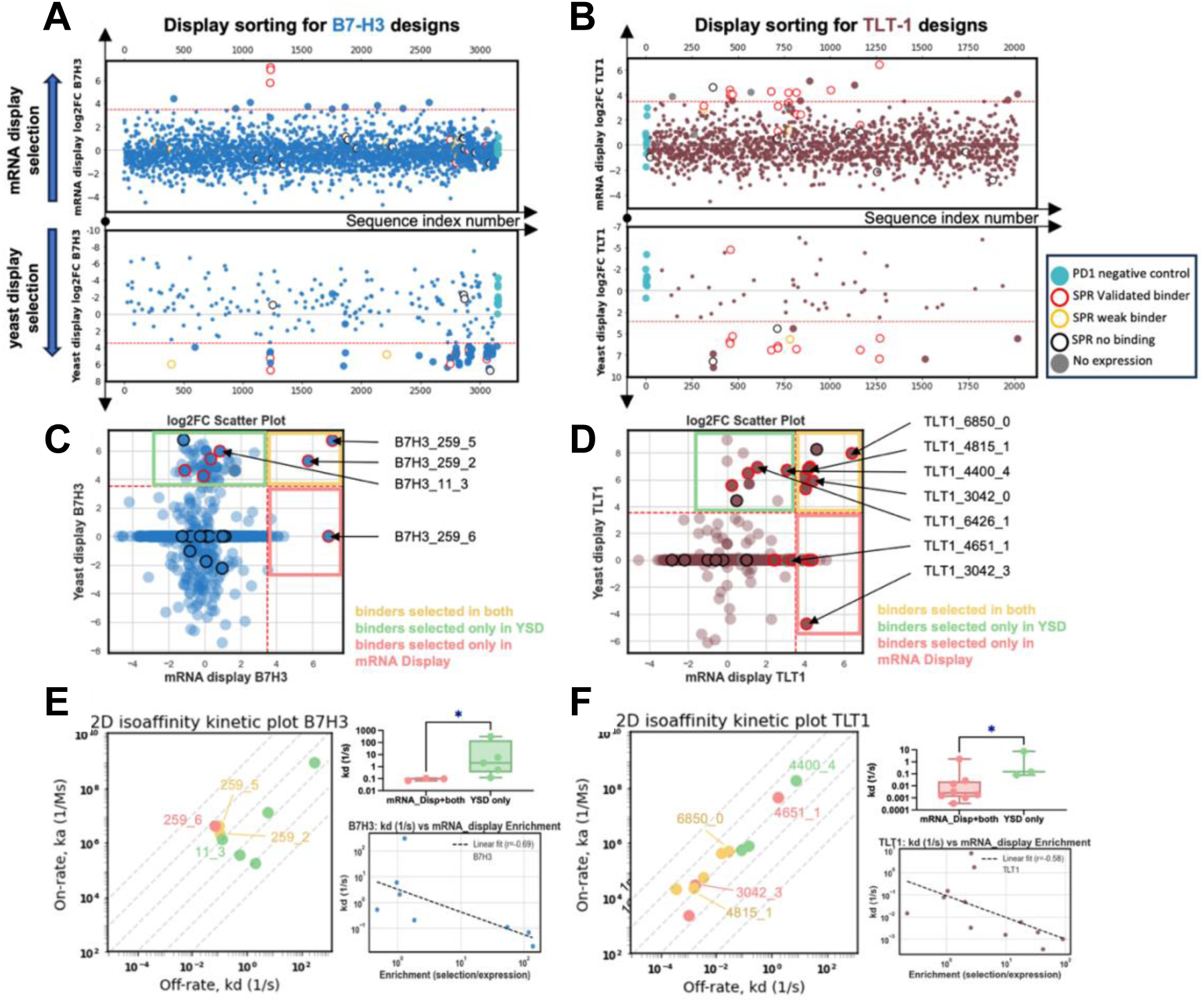
Parallel screening with mRNA and yeast display reveals enrichment profiles, platform-specific recovery, and kinetic bias. (A, B) Manhattan plots of log₂ fold-change (log₂FC) enrichment for all designs screened via mRNA and yeast display for B7-H3 (blue) and TLT-1 (brown), respectively. Validated binders (circles) are color-coded by platform of identification (mRNA only, yeast only, or both), and PD1 controls are included as negative controls. (C, D) 2D scatter plots comparing log₂FC values from mRNA (x-axis) and yeast (y-axis) display. Binders enriched by both methods fall in the upper-right quadrant (yellow box), while platform-specific hits fall in the lower-right (mRNA only, red box) or upper-left (yeast only, green box) quadrants. (E, F) Isoaffinity contour plots with enrichment quadrant labeled according to 2D scatter plots. Fast-off-rate binders (e.g., B7H3_11_3 and TLT1_6426_1) are primarily identified by yeast display, while stable, slow-off-rate binders (e.g., B7H3_259_5, TLT1_3042_3) are preferentially enriched by mRNA display. Correlation plots between enrichment score and kinetic off rate for validated binders. A negative correlation indicates that mRNA display is more sensitive to off-rate, with slower-dissociating binders showing higher enrichment.

The SPR validation results demonstrated high accuracy for both screening approaches. For B7-H3, 3 designs above the mRNA display enrichment cutoff were validated—all were true binders (0% false positive rate). Of 9 designs above the yeast display enrichment cutoff, 8 showed binding (11% false positive rate). For TLT-1, 13 designs enriched by mRNA display were tested; 10 were confirmed binders, 2 did not express, and only 1 did not bind—yielding a 7% false positive rate. For yeast display, 13 TLT-1 binders were tested and 11 bound, giving a 15% false positive rate. These results indicate that both mRNA and yeast display are effective for selecting high-affinity miniprotein binders, each with low false-positive rates. mRNA display offered slightly higher specificity and faster turnaround, while yeast display provided complementary coverage, particularly for binders that may be better stabilized in a cellular context.

Binding specificity was evaluated using a heatmap of enrichment values across different targets from mRNA display screening. This analysis confirmed that B7H3_259_5, B7H3_259_2, TLT1_3042_3, and TLT1_4815_1 showed minimal cross-reactivity with unrelated targets (**Figure S8**). PD1 control binders did not show enrichment on B7-H3 or TLT-1 selections in yeast or mRNA display. These results confirm that our computational design and high-throughput screening pipeline successfully identified binders with high specificity.

### Divergence in binder recovery between mRNA and yeast display reveals platform-specific biases

We first assessed library coverage of the two screening platforms by next generation sequencing of the library prior to selection. While both mRNA display and yeast display showed expression-dependent biases, mRNA display showed greater overall coverage. Specifically, mRNA display detected 83% of TLT-1 binders and 86% of B7-H3 binders, compared with 59% and 63%, respectively, for yeast display (**Figure S3**). These results suggest that the cell-free translation system used in mRNA display more effectively captures designed minibinders, likely by rescuing sequences that fail to express or fold efficiently in a cellular context. Improved library coverage is particularly important for ML-driven design as it supports more efficient feedback into computational design cycles.

To investigate the selection preferences of the two platforms, we compared the log₂ fold-change (log₂FC) enrichment scores for each design using two-dimensional scatterplots (**Figure 2C, D**). Each axis represents enrichment in one method, allowing us to visualize shared and platform-specific hits across the screened library. are those enriched in both platforms (upper-right quadrant) and represent high-confidence functional designs. Several of these, including *B7H3_259_5*, *B7H3_259_2*, *TLT1_6850_0*, *TLT1_3042_0*, and *TLT1_4400_4*, were confirmed by SPR to have strong binding affinities (**Figure 3B, S4**). These findings support the use of dual enrichment as a reliable predictor of binder functionality.

**Figure 3.**
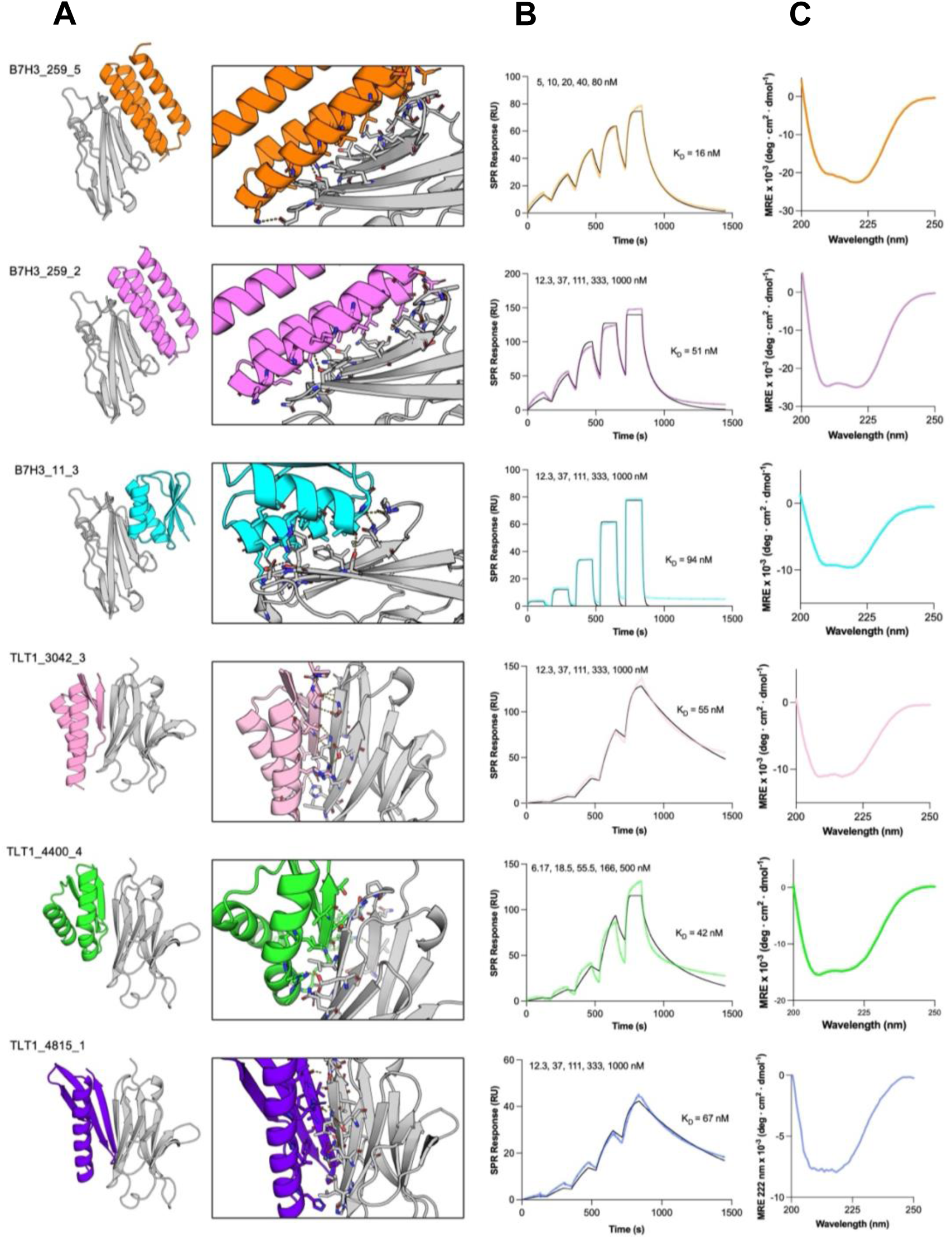
Structural modeling and biophysical validation of selected miniprotein binders. (A) Alphafold2 models of miniprotein binders (colored) in complex with their target (grey). (B) SPR binding affinity measurements. Solid black lines represent fits using a 1:1 kinetic model, with the dissociation constants derived from these fits. Analyte concentrations are shown above each plot. RU, response units. (C) Circular dichroism (CD) spectra of miniprotein binders at 25 °C.

Binders enriched only in mRNA display (lower-right quadrant) included validated hits such as *B7H3_259_6* and *TLT1_3042_3*. Although these designs exhibited high-affinity binding by SPR, they were not detected in yeast selection. These binders showed poor surface expression in yeast, suggesting that mRNA display may rescue functional designs that are penalized in cell-based systems due to instability or misfolding. Conversely, (upper-left quadrant) included *B7H3_11_3* and *TLT1_6426_1*. These designs demonstrated moderate affinity but fast dissociation rates, as seen in isoaffinity kinetic plots (**Figure 2E, F**).

The ten PD-1 control binders showed strong enrichment to PD-1 in yeast display selections. However, PD-1 binders with faster dissociation rates, such as AiD7, AiD15, and AiD19, were not enriched by mRNA display, reinforcing the presence of platform-specific kinetic bias across targets (**Figure S6**). This behavior likely reflects the differences in selection conditions between the two platforms. Our YSD display protocol employed two rounds of selection. In the first round of selection the biotinylated target proteins were premixed with streptavidin. As streptavidin is a tetramer, this allows for multivalent binding between multiple binders on the yeast surface (typically 10^4^-10^5^ copies per cell) ^20 21^ and target proteins linked to the same streptavidin molecule. Multivalent binding should stabilize weaker interactions and interactions with faster dissociation kinetics because both “arms” of an interaction need to dissociate simultaneously for the molecule to leave the cell surface. In the second round of YSD selection the target proteins were first mixed with the yeast cells and then a single wash was employed before adding fluorescently labeled streptavidin. During this second round of selection avidity effects should be less prevalent, but once streptavidin is added to the system multivalent interactions become possible. Together, the complementary selection biases between the two platforms highlights the value of integrating multiple screening technologies for comprehensive evaluation of ML-designed miniproteins.

### Biophysical validation of selected binders

A subset of binders enriched by mRNA display, yeast display, or both platforms were expressed in *E. coli* and characterized by surface plasmon resonance (SPR), size-exclusion chromatography (SEC), and circular dichroism (CD).

B7H3_259_5, one of top-enriched binders identified by both mRNA and yeast display, was predicted by AlphaFold2 to adopt a compact α-helical bundle structure interacting with residues I284, L293, and T298, on the B7-H3 β-sheet and residue F347 on a surface-exposed loop (**Figure 3A**)—previously reported to be involved in the immune inhibitory function of B7-H3 ^16^. Additional polar contacts involved residues A279, N282, T290, S343, R345, and D346. SPR confirmed high-affinity binding with an equilibrium dissociation constant K_D_ of ∼16 nM (**Figure 3B**). Size exclusion chromatography (SEC) showed a monomeric, non-aggregating species (**Figure S5A**). CD spectra were consistent with a well-folded α-helical structure, and thermal unfolding experiments showed no detectable unfolding up to 95 °C (**Figure S5B**).

B7H3_259_2 was designed from the same RFdiffusion scaffold as B7H3_259_5 and was also highly enriched by both selection platforms. The dissociation constant of this binder was determined to be ∼ 51 nM by SPR (**Figure 3B**). CD showed that the binder is folded correctly and showed no unfolding up to 95 °C (**Figure S3B**). SEC showed no evidence of aggregation (**Figure S5A**).

B7H3_11_3, which was enriched exclusively by yeast display, was predicted to adopt a mixed α/β fold with hydrophobic and polar contacts mediated by the two alpha helices on the binder and residues I284, Q286, F341, and F347 on B7-H3. SPR analysis revealed a moderate affinity (K_D_ ∼ 94 nM) and a fast dissociation rate (**Figure 3B**), consistent with its enrichment in yeast display but not mRNA display.

Among TLT-1 binders, TLT1_3042_3, enriched by mRNA display, was modeled as an α/β scaffold with an α-β-β-α-β topology. The final β-strand (residues 56–62) was predicted to pair with a β-strand on the TLT-1 target (residues 32–37), and residues 5, 15, 19, and 22 on the alpha helix are interacting with residues on the beta sheet surface of TLT-1 (**Figure 3A**). SEC confirmed that the binder was monomeric and non-aggregating (**Figure 3C**). SPR revealed a K_D_ of ∼ 55 nM (**Figure 3B**). The binder was thermostable, with a melting temperature (T_M_) around 87 °C (**Figure S5B**).

TLT1_4400_4, identified by yeast display, exhibited an α/β architecture (α-β-β-α-β topology), with the terminal β-strand (residues 57-62) predicted to pair with TLT-1 residues 32-37 (**Figure 3A**). This binder demonstrated strong affinity (K_D_ ≈ 42 nM), showed no aggregation on SEC (**Figure 3B**) and was thermostable up to 95 °C (**Figure S5B**).

TLT1_4815_1, identified by mRNA and yeast display, is predicted to adopt β-β-β-α fold, with predicted contacts at residues 10, 14, and 21 on the a-helix and residues 52-58 on the first β-strand, which pairs with residues 32-37 on TLT-1 (**Figure 3A**). Compared to the other binders, this design showed lower thermal stability (T_M_ ≈ 55 °C, onset ≈ 45 °C) and moderate SEC aggregation, consistent with the reduced stability of β-rich scaffolds.

### Evaluation of computational metrics for predicting experimental binding outcomes

Next, we evaluated the predictive value of several interface quality metrics using high-throughput screening outcomes from both mRNA and yeast display. Designs with log₂ fold-change enrichment greater than 3.5 (i.e., FC > ∼11.3) in either platform were designated as “binders,” while the remaining designs were treated as “non-binders.” We assessed multiple metrics derived from AlphaFold2 and RFdiffusion models, including PAE, interface pLDDT, and structural RMSD. These metrics were benchmarked against the experimental outcomes using receiver operating characteristic (ROC) curve analysis (**Figure 4A, B**).

**Figure 4.**
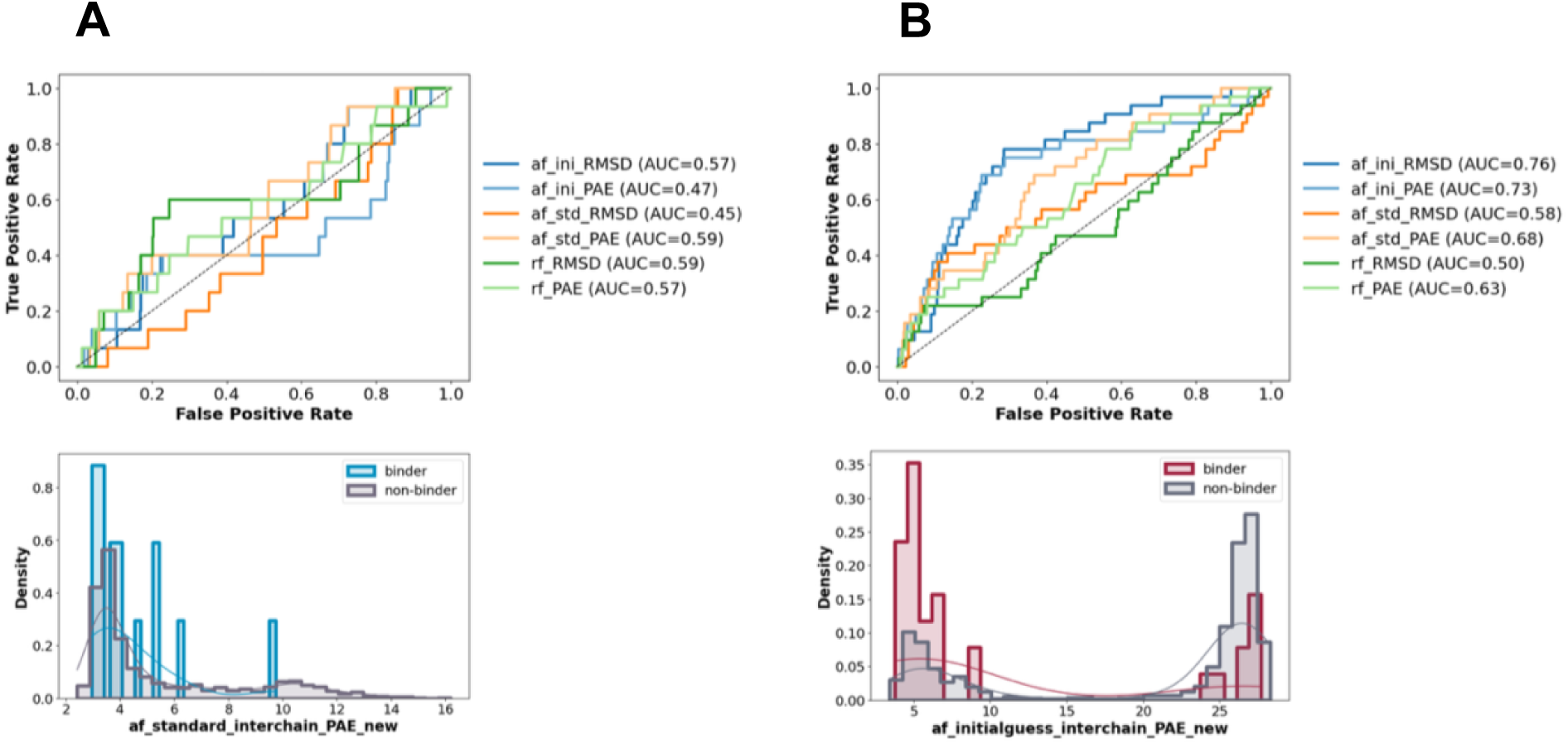
Predictive value of computational metrics for identifying binders. (A, B) ROC curves showing how well different computational metrics predict binders for B7-H3 and TLT-1. Metrics shows better predictivity for TLT-1 than B7-H3. (C, D) Histogram plots comparing metric values between binders and non-binders. Clear separation is seen for TLT-1, but not B7-H3, suggesting that prediction performance depends on the target.

The predictive power of individual metrics varied substantially between targets. For B7-H3, none of the metrics showed strong discriminative power. The best-performing metric — AlphaFold2 interface PAE—reached a modest area under the curve (AUC) of 0.59, indicating limited ability to distinguish binders from non-binders. In contrast, for TLT-1, several metrics showed promising predictive value. Notably, the interface PAE from AlphaFold2’s initial model guess achieved the highest performance, with an AUC of 0.73, suggesting that early-stage structure prediction may better capture useful interface features for this target.

Histogram distributions of selected metrics (**Figure 4C, D**) further illustrated partial separation between binder and non-binder populations for TLT-1, whereas little separation was observed for B7-H3. Together, these results underscore that, despite advances in structure prediction, reliable identification of binders using computational metrics alone remains challenging for certain targets, reinforcing the need for higher-throughput experimental feedback.

## DISCUSSION

In this study, we present a pipeline for miniprotein binder design that combines deep learning–based design tools with faster high-throughput experimental screening. ML-based scaffold generation and structure-guided sequence design enables more efficient sequence sampling in broader sequence space compared to traditional directed evolution methods using random sequence libraries, degenerate codons or site saturation mutagenesis. High-throughput screening provides sequence-to-function exploration at scale, efficiently identifying high-affinity binders among computational designs, and providing the insight for predictivity of interface quality metrics. This pipeline demonstrates a scalable, evolution-inspired framework in which machine learning-guided design is seamlessly integrated with high-throughput selection.

A central finding of this work is that mRNA display offers practical and mechanistic advantages for large-scale screening of ML-designed miniproteins. Its fully *in vitro* workflow enables shorter selection cycles and reduces experimental burden, making it particularly well suited for integration into iterative computational design pipelines. By systematically comparing mRNA display with yeast display using a shared oligonucleotide library, we demonstrate that mRNA display enables rapid and sensitive enrichment of functional binders, while exhibiting a bias toward slow-dissociating interactions. Because many therapeutic applications prioritize binders with slow dissociation rates, this feature makes mRNA display a suitable screening platform for therapeutic binder discovery.

Our results also demonstrate the value of high-throughput screening as a benchmark for assessing the predictive power of computational design metrics. While structure-based quality metrics provide modest predictive power overall, their performance was strongly target dependent. AlphaFold2 initial-guess models showed improved discriminative ability for TLT-1 binders, suggesting that early-stage structure predictions may encode informative features for certain targets. These findings underscore the limitations of relying on computational metrics alone for binder identification and emphasize the importance of large experimental datasets for improving ML-based scoring functions.

With a single round of computational design followed by high-throughput screening, we generated a panel of high stability and high affinity binders as therapeutic binder candidates. Overall, this work highlights the potential of integrating deep generative models with rapid display technologies to accelerate binder discovery. The flexibility and throughput of this platform support future application to other structurally challenging targets and therapeutic modalities.

## AUTHOR CONTRIBUTIONS

B.K., M.M., and Z.Y. conceived and designed the study. M.M. performed computational design of TLT-1 binders. Z.Y. performed computational design of B7-H3 binders. A.H. performed mRNA display selections. J.L. performed yeast display selections. M.M and Z.Y. conducted experimental validation of miniprotein binders and interpreted the data. T.K provided intellectual contributions for library construction. H.D. and A.N. assisted with analysis of computational metrics. M.M. and Z.Y. prepared the figures and wrote the manuscript with input from all authors. B.K. and A.B. supervised the project.

## METHODS

### Target Protein Modeling and Expression

The extracellular domains of human - (CD276) (UniProt Q5ZPR3: Gly 27 - Thr 461) and TLT-1 (TREM-like transcript-1) (UniProt Q86YW5: Gly20 – Ply125) were selected as regions to include as binder targets. Human B7-H3 structure was modeled using AlphaFold2 to obtain the target protein structure for design. Human TLT-1 crystal structure (PDB 2FRG) was used in the design as target protein structure ^17^.

For downstream display and binding assays, the target protein was expressed in mammalian cells and biotinylated with Avitag^TM^. Human B7-H3 expressed in human 293 cells (HEK293) was purchased from Acro Biosystem (Acro, Biotinylated Human B7-H3 (4Ig), Cat. #B7B-H82E8). Human TLT-1 was expressed in-house in EXPI293F cells (Thermo Fisher Scientific, Cat. # A14527) with a human serum albumin signal peptide for secretion to the culture medium. Plasmid DNA with C-terminal polyhistidine tag and Avitag^TM^ was synthesized and cloned (Twist biology), transiently transfected into EXPI293F and expressed according to the manufacturer’s protocol. EXPI293F cells were maintained at 37°C with 8% CO2 at 250 rpm without antibiotics and were harvested 96 hours after transfection. The supernatants were separated from the cells by spinning at 3000 rcf (4°C) and passing through 0.4µm filters. The supernatants were purified with penta-Ni resin (Marvelgent) according to the manufacturer’s protocol. BirA biotin-protein ligase was used to biotinylate the Avitag^TM^ using the manufacturer’s protocol (Avidity, BirA biotin-protein ligase standard reaction kit, BirA500)

### Computational scaffold generation, sequence design and filtering based on computational metrics

*De novo* miniprotein binders were designed using a deep learning–based pipeline. First, structural scaffolds were generated using Rfdiffusion ^4^. For each target, B7-H3 and TLT-1, interface “hotspot” residues (2–4 residues per input) were manually selected based on exposed surface accessibility and target epitope relevance (**Figure 1**). These residues were used to guide the model toward generating binders that interact with the defined target interface. Each RFdiffusion cycle produced ∼10,000–20,000 backbone scaffolds. Initial filtering was applied using radius of gyration and contact metrics (via BioPython), retaining ∼1,250 scaffolds per target for downstream sequence design. The generated backbone structures were then subjected to sequence design using ProteinMPNN. Between 4-7 ProteinMPNN sequences were generated per backbone, resulting in ∼8000 designed miniprotein binder sequences.

These designed sequences were evaluated by three independent structural prediction methods: AF2 ^2^, AF2 initial guess ^9^, and RoseTTAFold (RF) ^18^. The pLDDT of the designed miniprotein binders were used to evaluate the binder quality. The binders with pLDDT > 80 were selected for interface quality evaluation. Structural confidence of the designed binder interface, interface PAE and RMSD to the scaffold pose to evaluate binder-target interaction quality. Binders with PAE < 10 and RMSD < 5 in any of structure prediction methods were selected for experimental screening. The final experimental library consisted of 2,009 TLT-1-targeting and 3,149 B7-H3-targeting sequences (**Figure 1a**).

### Oligo Library Construction

Selected DNA sequences were codon-optimized for expression in *E. coli* and synthesized as an oligonucleotide pool. The oligo pool was amplified by PCR with overhang primers for yeast display or mRNA display. mRNA overhang primers added the RBD region and puromycin linking region to each DNA sequence. Final DNA libraries for mRNA display were verified with Azenta Amplicon EZ sequencing. Yeast display overhangs added the overlapping region to the pCTcon2 vector for yeast homologous recombination. DNA inserts were prepared according to a standard protocol and co-transformed with linear pCTcon2 vector to EBY100 yeast cells ^13^. Plasmid DNA was extracted from Yeast libraries and PCR prepared for Amplicon EZ sequencing.

### Yeast display screening

#### Yeast cell growth and protein expression

EBY100 yeast cells co-transformed with linear pCTcon2 vector and synthesized DNA library were inoculated into 5ml SDCAA (20g/L Dextrose, 6.7g/L Yeast Nitrogen Base, 5g/L Casamino Acids, 8.6g NaH_2_PO_4_·H_2_O, 5.4g/L Na_2_HPO_4_, pH4.5) ^21^ and grown by shaking at 250rpm and 30 °C for 16 hours. Overnight (O/N) cell culture was then inoculated into 5ml SDCAA medium at a cell density of 1×10^7 cells/ml. This initial P1 yeast library was sent for Amplicon EZ sequencing with Azenta and used as the reference library for sequencing data analysis. The subculture of initial P1 yeast library was then grown at 30 °C until the OD₆₀₀ reached 2.0. A total of 5 × 10⁷ cells were harvested by centrifugation at 2000 × g for 3 minutes, and the resulting cell pellet was resuspended in 5 mL SGCAA medium (20g/L Galactose, 6.7g/L Yeast Nitrogen Base, 5g/L Casamino Acids, 8.6g NaH_2_PO_4_·H_2_O, 5.4g/L Na_2_HPO_4_, pH6.0) and cultivated at 20 °C to induce protein expression for 16-18h.

#### Fluorescence-Activated Cell Sorting (FACS)

For sorting, 1×10^7^ cells were stained with 50μl of mixture containing 2μM biotinylated target protein (PD-1, TLT-1 and B7-H3) premixed with 500nM (150 ug/mL) R-Phycoerythrin-Conjugated Streptavidin (SAPE) (Thermo Fisher Scientific, Catalog #S866) and 5μg/ml anti-c-Myc Alexa 488 (Biolegend, Catalog#626812). Cells were washed once with 1 mL PBS+0.1% BSA prior to FACS. Cells displaying double-positive signals for PE and Alexa Fluor 488 were sorted and subsequently grown in 5 mL SDCAA medium at 30 °C, followed by expression in SGCAA as described above. For the second sorting, 1 × 10⁷ cells were stained with 50 μL of 200 nM biotinylated target protein and 4 μg/mL chicken anti-c-Myc antibody (Thermo Fisher Scientific, Catalog #A-21281), washed once with 1 mL PBS+0.1% BSA, and stained with 1 μg/mL SAPE and 2 μg/mL goat anti-chicken IgY (H+L) secondary antibody (Invitrogen, Catalog #A11039).

#### Plasmid extraction and PCR for next-generation sequencing (NGS)

Plasmid DNA was extracted from the yeast cell pellet obtained from 1 mL of SDCAA culture of sorted cells using the Zymoprep™ Yeast Plasmid Miniprep II Kit (Zymo Research, Catalog #D2004). Selected miniprotein variant genes were then amplified using Q5 High-Fidelity DNA Polymerase with the forward primer:

Seq_pCTct20_fwd (5′-CAATAGCTCGACGATTGAAGGTAG-3′)

and the reverse primer:

Seq_pCTct20_rev (5′-ACACTGTTGTTATCAGATCTCG-3′)

The PCR products were purified using the QIAquick PCR Purification Kit (QIAGEN, Catalog #28104). DNA was diluted to 20 ng/mL and sent to Azenta for Amplicon EZ sequencing.

### mRNA display screening

#### RNA library preparation, Transcription

A 1 mL transcription reaction (adapted from NEB T7 RNA polymerase protocols) containing amplified DNA (100 μL), 1x T7 RNA Polymerase Buffer, 1 mM Dithiothreitol (DTT), 16.5 mM MgCl_2_, 5 mM rNTPs, and 5 units/μL T7 RNA Polymerase (NEB M0251) was incubated overnight at 37 °C. Successful transcription reactions appear cloudy due to the precipitation of magnesium pyrophosphate. Then, 115 μL of 10x DNAse I buffer and 30 μL of DNAse I (Promega PR-M6101) was added to the solution and incubated at 37 °C for 1h. After DNAse incubation, EDTA, NaCl, and 100% isopropyl alcohol were added to final concentrations of 37.5 mM, 150 mM, 45%, respectively. The solution was then centrifuged at 14,000 rpm for 15 mins to pellet the RNA. The supernatant was removed, and the pellet was washed with excess 70% ethanol. This solution was briefly centrifuged at 14,000 rpm, the supernatant was removed, and the RNA pellet was allowed to dry at RT. The RNA pellet was then solubilized in 100 μL of MQ-H_2_O and an equal volume (100 μL) of 2x RNA loading dye. The sample was heated at 95° C for 7 mins, then run on a large scale 12% Urea-PAGE gel at 230 V for 1.5 h in a Tris-Borate-EDTA (TBE) buffer. The desired RNA band was then visualized under UV illumination at 254 nm on a silica-coated thin-layer chromatography plate and excised from the gel. The excised band was crushed into fine pieces, and the RNA was extracted with 0.3 M NaCl (2 x 1 h incubation at RT). To collect the RNA, the gel was pelleted by centrifugation at 14,000 rpm and the supernatant isolated. The supernatant was then passed through a 0.45 μm filter before addition of 2x volume of 100% ethanol. The solution was mixed vigorously, then was centrifuged at 14,000 rpm for 15 mins to pellet the RNA. The pellet was washed with excess 70% ethanol, briefly centrifuged at 4,000 rpm, and then allowed to dry at RT. The RNA was solubilized in MQ-H_2_O and the concentration was determined by nanodrop and adjusted as necessary. Purity was confirmed via an analytical scale 12% Urea-PAGE gel. RNA was stored at −80° C until use.

#### RNA library preparation, Puromycin Linking

RNA of single sequences or randomized libraries were covalently linked to puromycin via an adapted Y-ligation method. A solution with 1 μM RNA, 20% DMSO, 1.5 μM puromycin linker (P29), 1x T4 RNA ligase buffer, and 1 unit/μL T4 RNA ligase I (PR-M1051) in MQ-H_2_O was incubated at 37° C for 0.5 h. Then, 1x volume of precipitation solution (0.6 M NaCl, and 50 mM EDTA), 0.02x volume of 5 mg/mL glycogen, and 2x volume 100% ethanol were added. This mixture was briefly vortexed, then centrifuged at 13,000 rpm for 15 mins to pellet the puromycin-linked RNA. The supernatant was removed, washed with excess 70% ethanol, and briefly centrifuged at 16,000 x g. The supernatant was removed, and the pellet was allowed to dry at RT. Once dry, the puromycin-linked RNA pellet was reconstituted in 1/10th the volume of the p-linking reaction (10 μM RNA). The efficiency of the reaction was determined by 12% Urea-PAGE Gel (National Diagnostics, EC-833).

#### Translation and Reverse Transcription

A 2.5 uL translation of puromycin-linked RNA was performed using a custom NEB PURExpress® kit (-aa, -tRNA, -RF123) -E6850Z for each target protein. Translations were incubated at 37° C for 30 mins, followed by a 10 min incubation at RT to facilitate fusion of peptide to its mRNA strand. EDTA was then added to a final concentration of 17 mM to dissociate the ribosome, and the mixture was incubated at 37°C for 20 mins. Then, complementary DNA was appended by a reverse transcription reaction containing all the translation product, 0.6 mM dNTPs, 5 μM reverse primer, 62.5 mM Tris-HCl pH 8.3, 37.5 mM Mg(OAc)2, 25 mM KOH, 2.5 x M-MLV reverse transcriptase H (-) point mutant (Promega, M3681, supplied at 40x), and MQ-H_2_O (to reach final volume). This solution was incubated at 42° C for 1h.

#### HA Purification

Fusions containing an HA peptide sequence were diluted by 10x with TBS-T and added to anti-HA magnetic beads (ThermoFisher catalog number: 88836) at a 4:1 ratio of bead slurry to initial IVT volume. The bead slurry rotated at RT for 45 min, after which beads were washed 3x with TBS-T. 25 uL (10X IVT volume) of 2 mg/mL HA peptide in selection buffer (1X phosphate buffered saline, 0.05% Tween-20) was then added to the beads, and fusions were eluted by rotation at RT for 1 h.

#### Target protein Immobilization onto Streptavidin Beads

Biotinylated target protein was immobilized onto Dynabeads^TM^ M-280 Streptavidin beads (ThermoFisher, catalog #11205D). Excess target protein in selection buffer was added to streptavidin beads such that, when combined with the volume of HA eluted fusions, enough protein was immobilized to arrive at a final concentration of 200 nM in solution. A separate streptavidin immobilization assay was performed prior to this to determine the quantity of beads needed to immobilize this concentration of each protein. Immobilization occurred at 4 °C for 30 mins. Then, free biotin in H_2_O was added to a final concentration of 25 μM to occupy any remaining biotin binding sites. The solution rotated for an additional 15 mins at 4 °C to facilitate biotin blocking. Once complete, protein-immobilized SA beads were washed 3x with 200 uL selection buffer.

#### Selection and cDNA Elution

25 uL of the eluted HA eluted fusions were added to the protein immobilized SA beads. The libraries were incubated for 1 h at 4 °C. After incubation, the supernatant was removed, and protein immobilized-SA beads were washed 3x with 200 uL selection buffer. Protein-SA beads were then resuspended in 25 μL of H_2_O and heated at 95° C for 5 mins to elute cDNA. The supernatant containing the eluted cDNA was recovered and analyzed via qPCR.

#### qPCR validation for selection result

qPCR was performed with an Applied Biosystems Viia7 qPCR Real-Time qPCR Instrument. Each qPCR sample contained 0.25 μM forward and reverse primers and 1 μL of experimental sample in 1x SsoAdvanced Universal SYBR Green Supermix (Bio-Rad, 172-5271). qPCR standards were prepared by reverse transcription of a known quantity of RNA into cDNA (assuming 100% yield) and prepared in 10-fold serial dilutions to 2e9, 2e8, 2e7, 2e6, 2e5, and 2e4 molecules. For amplification, initial heating began at 50 °C for 2 min followed by 10 min 95 °C incubation. Cycling then occurred between 95 °C for 15 sec and 60 °C for 1 min, with cycler heating acceleration held at 1.6 °C between each step. A total of 40 cycles were performed in each qPCR run. During each elongation step, SYBR green fluorescence was measured with ROX as a passive reference. Following the run, standard curves were generated and used to calculate cDNA quantities.

#### PCR Amplification and NGS

Amplification of recovered cDNA was performed until bands could be observed via DNA gel, usually 15-30 cycles. After DNA purification with a GeneJET PCR Purification kit (ThermoFisher, Catalog # K0702), DNA was diluted to 20 ng/mL and sent to Azenta for Amplicon EZ sequencing.

### Next-Generation Sequencing and Data Analysis

Raw sequencing reads were obtained in FASTQ format from Azenta Amplicon EZ sequencing. Quality control and adapter trimming were performed using fastp ^22^, with default parameters unless otherwise specified. Low-quality bases and reads with average Phred quality scores below Q30 or read lengths shorter than 170 bps were discarded.

Filtered reads were then processed using an in-house Python script to identify sequences containing the expected primer-flanking regions. Reads matching the correct pattern were retained, and the core coding region was extracted and translated into amino acid sequences. Sequences with frameshifts or premature stop codons were excluded from downstream analysis.

The resulting peptide sequences were aligned to a reference library of designed variants using pyahocorasick ^23^ for fast and memory efficient library exact string search. A count table was generated to quantify the frequency of each unique sequence across different samples and replicates. Further downstream analyses were performed using custom Python scripts. The fold change (FC) was calculated based on the NGS result after selection against target protein and NGS result of reference library of mRNA display and Yeast display.

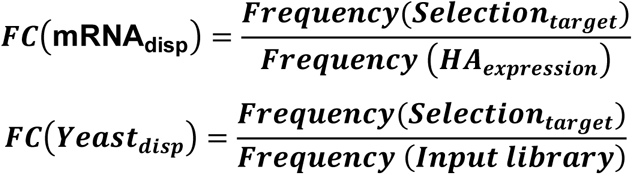

### Cloning, expression, and purification of Miniprotein expression for biochemical and biophysical validation

*Escherichia coli* BL21 cells (Thermo Fisher Scientific, USA) were used for protein expression due to their high efficiency and compatibility with T7 promoter-driven expression systems. The gene of Miniprotein binders was cloned into the expression vector pET-21a(+), containing an N-terminal His-tag and UV detection region. Plasmid DNA was transformed into competent BL21 cells using the heat-shock method. A single colony was picked and grown in 5 mL of LB medium with the Ampicillin overnight at 37°C, with shaking at 250 rpm. The overnight culture was used to inoculate 100ml of fresh TB culture. Cells were induced with IPTG (final concentration of 1 mM) at OD600 of 0.6-0.8. The temperature was then dropped to 16°C and cells were shaken at 250 rpm overnight. Cells were harvested by centrifugation at 4,000 x g for 30 minutes at 4°C. The cell pellet was frozen and stored at −80°C. Cells were lysed in lysis buffer (500mM NaCl, 50mM Tris HCl pH 7.5, 10mM Imidazole, PMSF 1mM, Pepstatin 10 µM, Bestatin 1 µM, Leupeptin 10 µM) by sonication on ice (5 seconds on, 5 seconds off, at 70% amplification for 5 minutes).

The lysate was clarified by centrifugation at 12,000 x g for 30 minutes at 4°C. The supernatant was subjected to batch binding with Ni-penta resin (Marvelgent Biosciences, Catalog #11-0228-010) under gentle shaking at 4°C for 30 minutes to allow protein binding. The column was washed with 20 column volumes of high salt wash buffer (1M NaCl, 50mM Tris HCl pH 7.5, 10mM Imidazole) and 2.5 column volumes of wash buffer (500mM NaCl, 50mM Tris HCl pH 7.5, 25mM Imidazole) to remove non-specifically bound proteins. The target protein was eluted with elution buffer (500mM NaCl, 50mM Tris HCl pH 7.5, 500mM Imidazole).

### SPR

Surface Plasmon Resonance (SPR) was used to validate the binding affinity between selected miniprotein binders and target protein. Biotinylated target protein (B7-H3 or TLT-1) was immobilized onto a NeutrAvidin™ sensor chip (Cytiva, Series S Sensor Chip NA, Cat. #29699622) at a concentration of 100 nM, yielding approximately 500 response units (RU). Miniprotein samples were prepared in 5 dilution series and injected over the chip surface using the Biacore™ 8K SPR system. The running buffer consisted of PBS with 0.05% Tween-20. Single-cycle kinetic experiments with five miniprotein concentrations were conducted with 120-second association phase and a 600-second dissociation phase, at a flow rate of 30 µL/min. The association rate constant (k_on_), dissociation rate constant (k_off_), and the equilibrium dissociation constant (K_D_) were determined by fitting to a 1:1 kinetic binding model within the Biacore™ Insight Evaluation Software.

### Assessment of Miniprotein secondary structure and stability with Circular dichroism (CD)

Each purified miniprotein binder was centrifuged at 14,000 × g for 10 min at 4°C and the supernatant diluted to 20 µM in PBS. Circular dichroism spectra and temperature melts were acquired with a Jasco CD J-815 C instrument. Thermal unfolding was monitored by ramping at 1°C intervals from 20°C to 90°C and measuring the CD signal at 222 nm during the temperature ramp. Molar ellipticity was calculated with the following expression: [θ]=θ/(10 × c × l) where l is the path length in cm. The midpoint for thermal unfolding was determined by fitting the data to the temperature dependent Gibbs-Helmhotz Equation using non-linear regression tools in Microsoft Excel.

### Size-exclusion chromatography (SEC)

Following nickel affinity chromatography, miniproteins were subjected to size-exclusion chromatography (SEC) analysis using a Superdex 75 Increase 10/300 GL column (Cytiva 29148721). Fractions corresponding to each peak were collected and analyzed by SDS-PAGE under reducing conditions to determine the composition of each fraction.

