## Supplementary data for "Enhancing ML-based binder design with high-throughput screening: a comparison of mRNA and yeast display technologies"

**Table of Contents:**

**Figure S1.** Computational filtering and distribution of interface quality metrics for designed miniprotein binders

**Figure S2.** t-SNE projection of binder ESMFold embeddings

**Figure S3.** Analysis of binder expression biases between mRNA display and yeast display

**Figure S4.** SPR traces of additional TLT-1 binders

**Figure S5.** Biophysical characterization of selected miniprotein binders

**Figure S6.** High-throughput screening of previously validated PD-1 miniprotein binders

**Figure S7.** SDS-PAGE analysis of purified miniprotein binders selected for biochemical and biophysical validation

**Figure S8.** Binding specificity matrix for miniprotein binders based on mRNA display enrichment

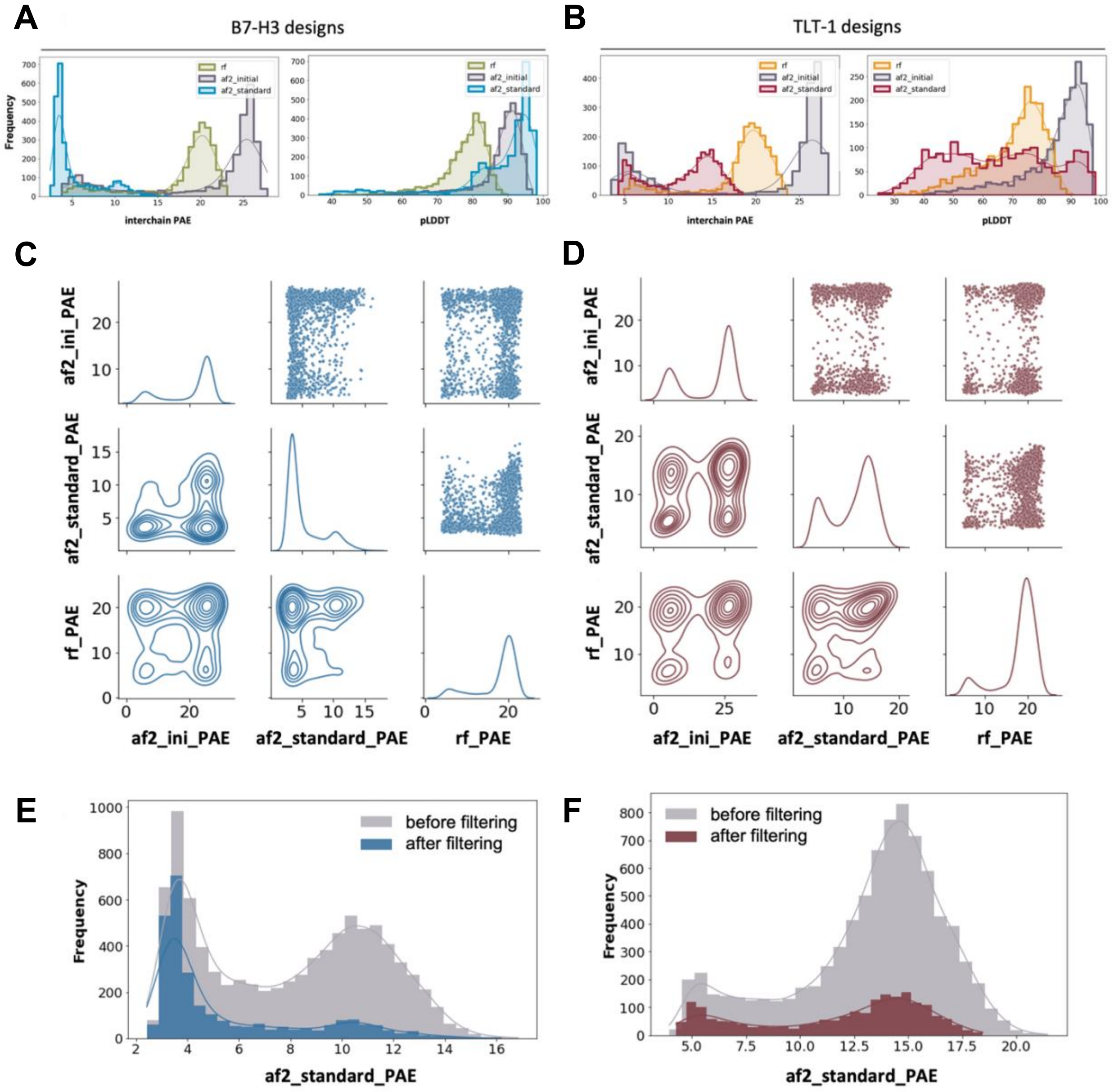


**Figure S1. Computational filtering and distribution of interface quality metrics for designed miniprotein binders.** (A, B) Distributions of individual structure prediction and interface metrics used to evaluate binder candidates for B7-H3 (A) and TLT-1 (B). Metrics include pLDDT and interface PAE. Each curve represents output from a different structure prediction model (AF2, AF2 initial guess, RF). (C, D) Pairwise correlation plots between computational metrics for B7-H3 and TLT1 designs, respectively, highlighting relationships between binder confidence and predicted interface quality. (E, F) Histogram comparison of raw (unfiltered) and filtered design libraries for both targets. Filtering retains metric diversity while enriching for candidates predicted to form high-quality interfaces.

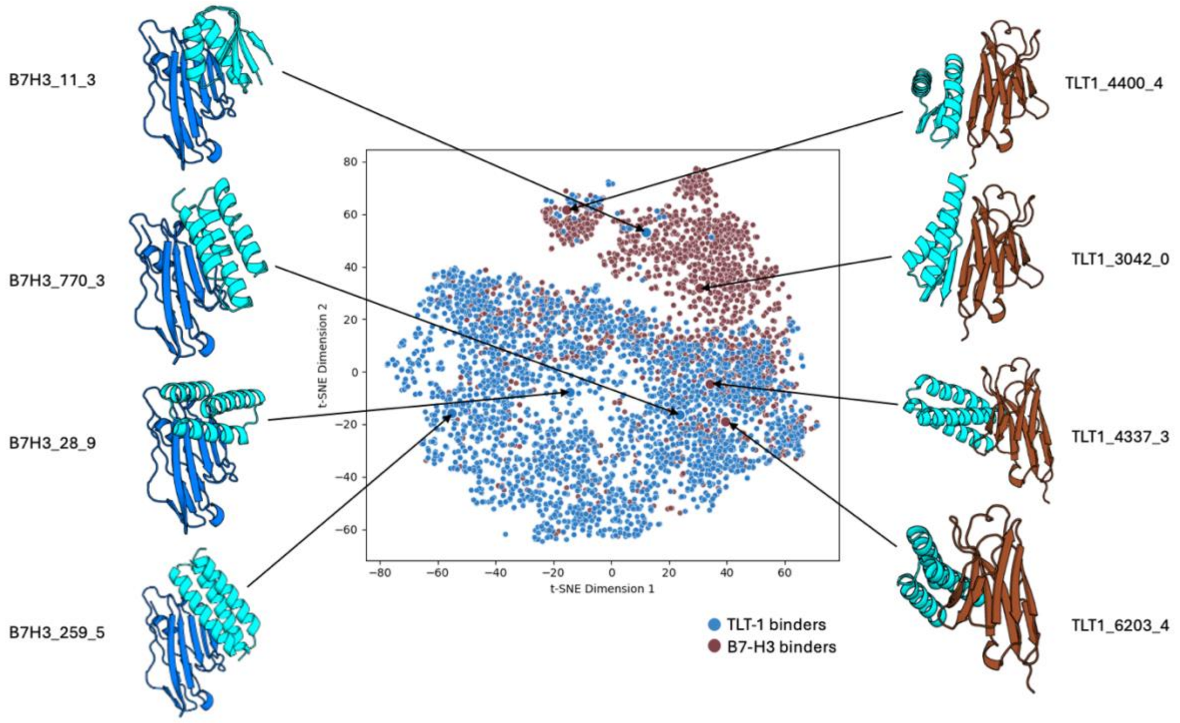


**Figure S2. t-SNE projection of binder ESMFold embeddings.** 2D t-SNE projection of the final experimental design library showing sequence diversity of selected binders. B7-H3 and TLT-1 designs occupy distinct regions in sequence space.

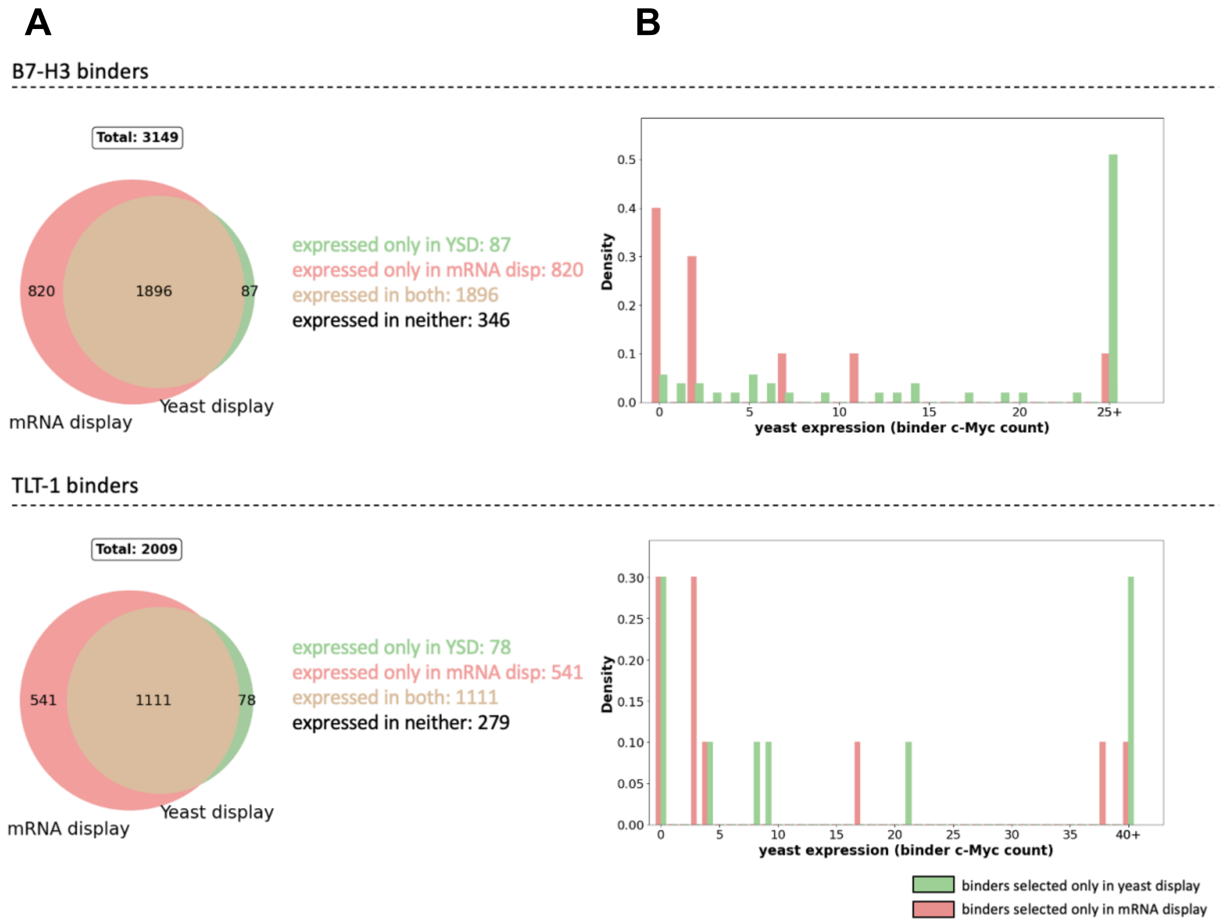


**Figure S3. Analysis of binder expression biases between mRNA display and yeast display.** (A) Venn diagram comparing expression coverage between mRNA display and yeast display for B7-H3 binders (top) and TLT-1 binders (bottom). For mRNA display, expression was measured by HA count of the library before selection; for yeast display, c-Myc count. mRNA display has greater expression coverage overall. Specifically, there are 820 B7-H3 binders and 541 TLT-1 binders expressed only in mRNA display but not detected in yeast. (B) Density histogram of yeast library c-MYC counts for TLT-1 binders selected by mRNA display only (red) or yeast display only (green). For binders observed only in mRNA display selections, many lack c-Myc counts, suggesting that expression issues likely explain why these binders were missed in yeast display.

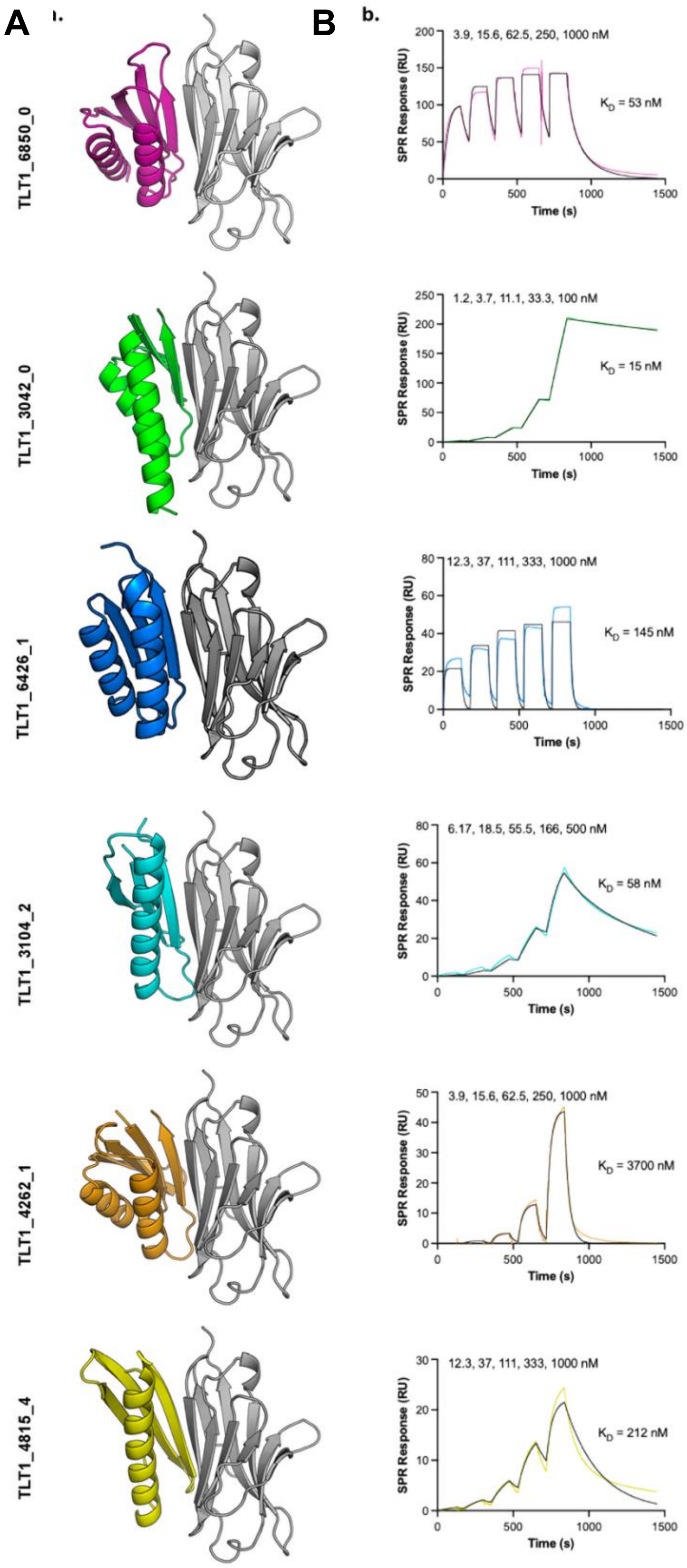
A
B
b.
TLT1_6850_0
3.9, 15.6, 62.5, 250, 1000 nM
KD = 53 nM
TLT1_3042_0
1.2, 3.7, 11.1, 33.3, 100 nM
KD = 15 nM
TLT1_6426_1
12.3, 37, 111, 333, 1000 nM
KD = 145 nM
TLT1_3104_2
6.17, 18.5, 55.5, 166, 500 nM
KD = 58 nM
TLT1_4262_1
3.9, 15.6, 62.5, 250, 1000 nM
KD = 3700 nM
TLT1_4815_4
12.3, 37, 111, 333, 1000 nM
KD = 212 nM
SPR Response (RU)
Time (s)

**Figure S4. SPR traces of additional TLT-1 binders.** (A) Alphafold2 models of miniprotein binders (colored) in complex with their target (grey). (B) Solid black lines represent fits using a 1:1 kinetic model, with the dissociation constants derived from these fits. Analyte concentrations are shown above each plot. RU, response units.

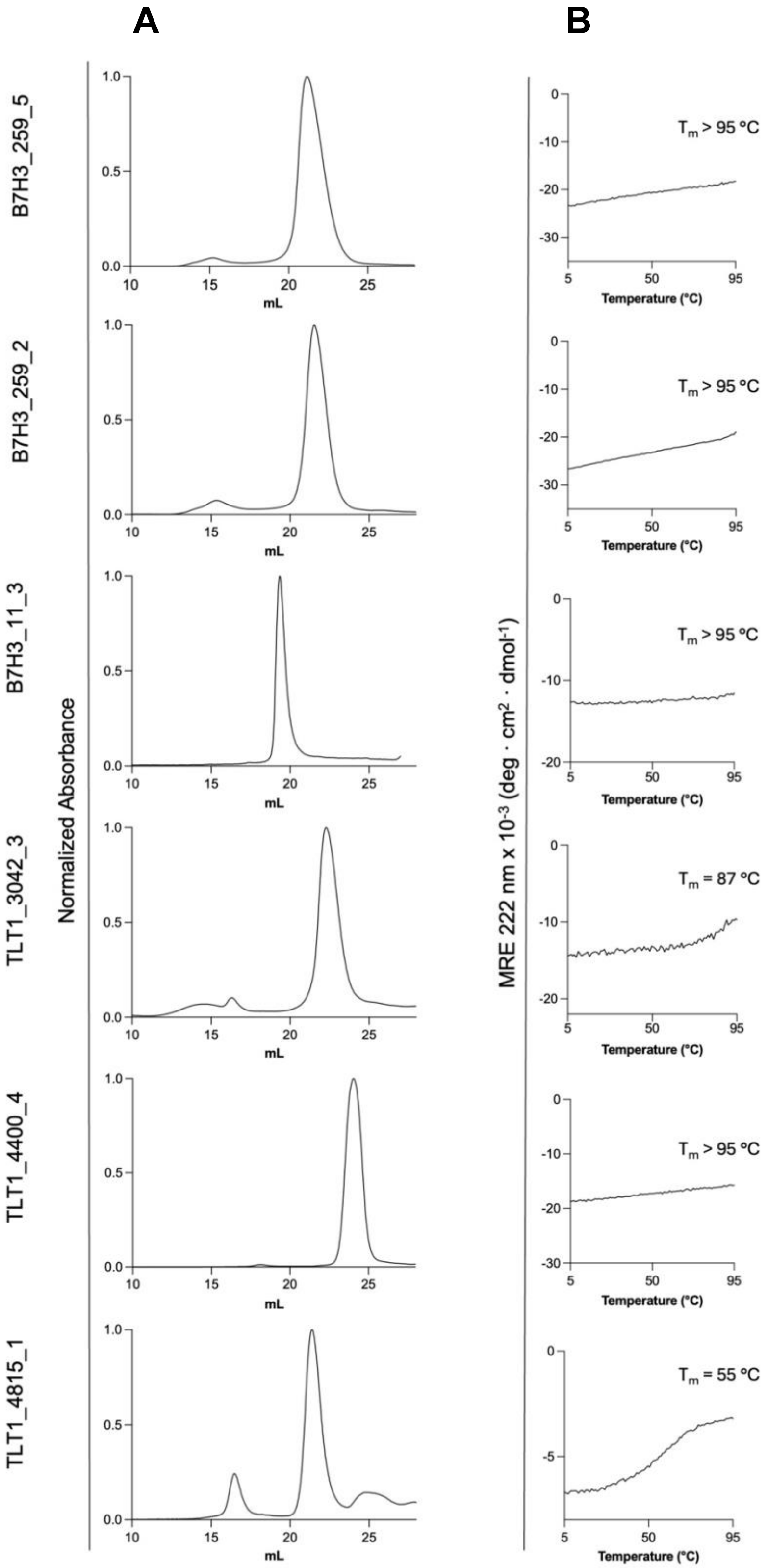
A
B
B7H3_259_5
B7H3_259_2
B7H3_11_3
TLT1_3042_3
TLT1_4400_4
TLT1_4815_1
Normalized Absorbance
mL
MRE 222 nm x 10-3 (deg · cm2 · dmol-1)
Temperature (°C)
Tm > 95 °C
Tm > 95 °C
Tm > 95 °C
Tm = 87 °C
Tm > 95 °C
Tm = 55 °C

**Figure S5**. **Biophysical characterization of selected miniprotein binders**. (A) Size exclusion chromatography traces of selected binders following purification by nickel affinity chromatography. (B) Circular dichroism signal at 222-nm wavelength as a function of temperature. Five of the six de novo designs show ultra-high thermostability ($T_m$ > 85 °C).

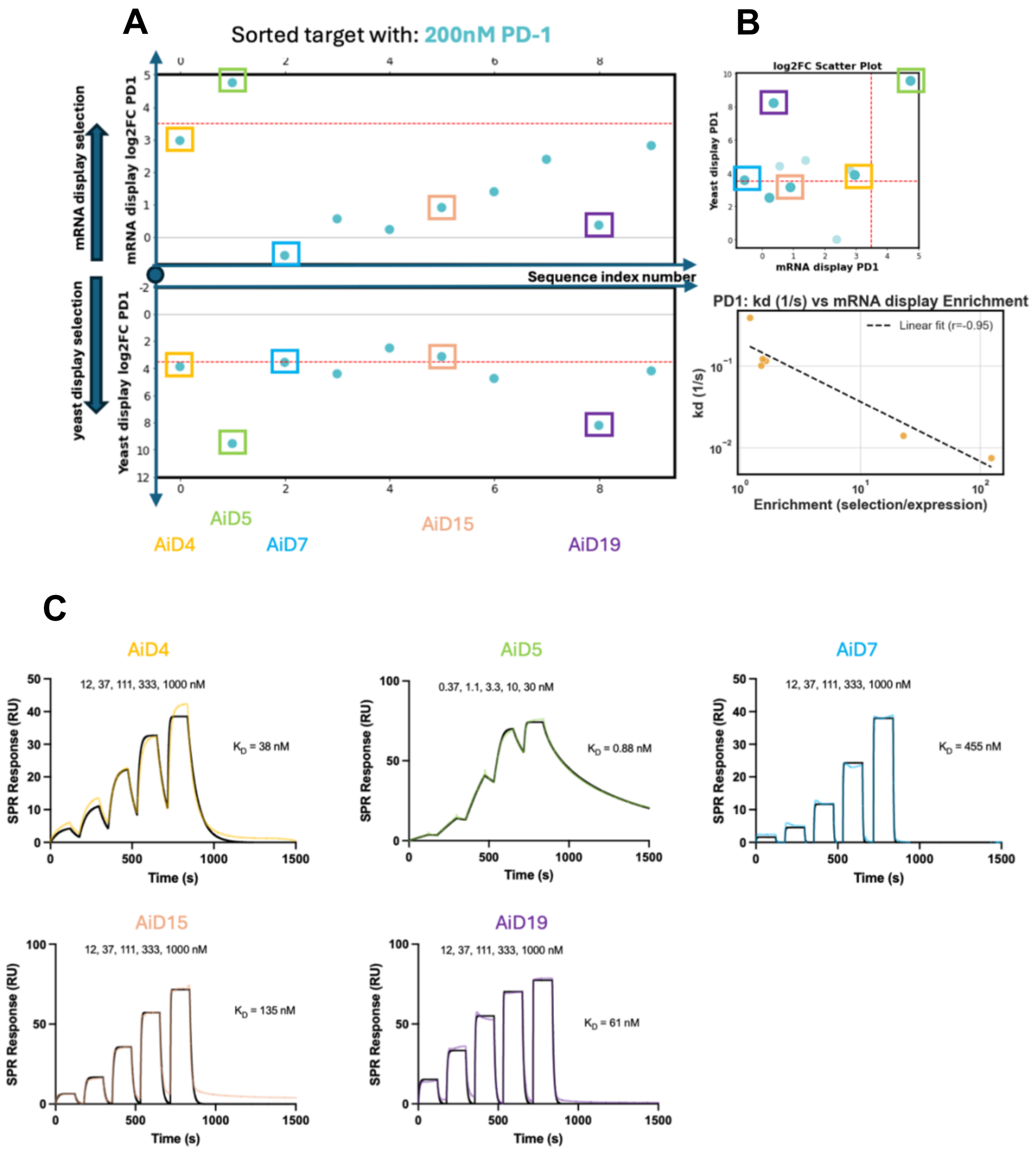

**Figure S6. High-throughput screening of previously validated PD-1 miniprotein binders.** Ten miniprotein binders previously confirmed to target PD-1 were spiked into the oligo library and screened against biotinylated PD-1 using both mRNA and yeast display platforms. (A) Manhattan plots showing enrichment ($log_2FC$) values for each design from mRNA display (top) and yeast display (bottom). (B) Scatter plot comparing $log_2FC$ values between the two platforms. Binders enriched in both platforms are located in the upper-right quadrant. Correlation plot between enrichment score and kinetic off rate. A negative correlation indicates that slower-dissociating binders show higher enrichment. (C) SPR traces of PD-1 binders previously validated with SPR [6]. Binders with faster dissociation rates, such as AiD7, AiD15, and AiD19, were preferentially enriched in yeast display, while binders with slower dissociations such as AiD4 and AiD5 were recovered by both platforms.

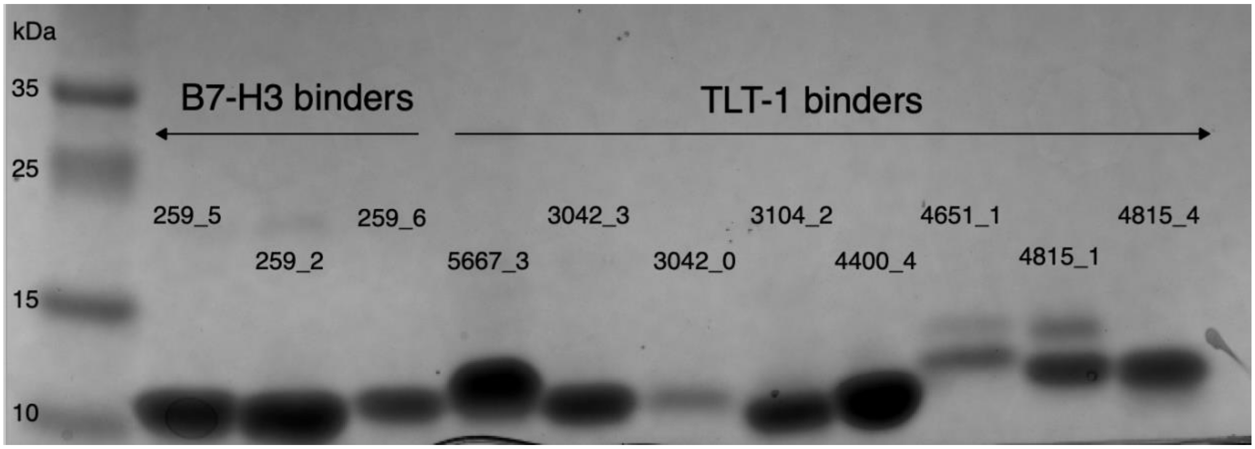


**Figure S7. SDS-PAGE analysis of purified miniprotein binders selected for biochemical and biophysical validation.** Representative Coomassie-stained SDS-PAGE analysis of B7-H3 or TLT-1 miniprotein binders selected for biophysical validation. Designs were expressed in *E. coli* and purified via affinity chromatography for downstream structural and functional studies. Lanes are labeled with design IDs, and corresponding apparent molecular weights are shown. Most purified proteins show a single band near the expected size (~10–15 kDa), indicating successful expression and high purity.

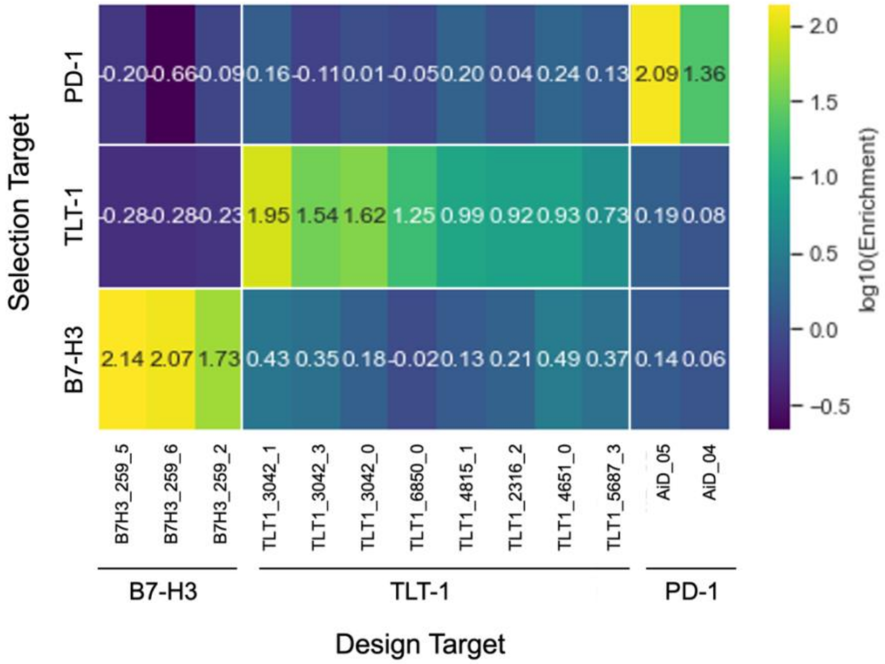


**Figure S8. Binding specificity matrix for miniprotein binders based on mRNA display enrichment.** Heatmap showing $\log_{10}$ enrichment values of selected binders screened against B7-H3, TLT1, and PD-1 targets using mRNA display. Rows indicate selection targets, and columns represent individual binders. High off-target enrichment suggests potential non-specific binding.
